# Ethical Considerations of Mitigating Data Loss: VLADISLAV, a Manifesto for Reliable Home Cage Systems

**DOI:** 10.64898/2026.01.20.700603

**Authors:** Davor Virag, Ana-Marija Virag, Jan Homolak, Pia Kahnau, Ana Babić Perhoč, Antonia Krsnik, Luka Mihalic, Ana Knezović, Jelena Osmanović Barilar, Mario Cifrek, Vladimir Trkulja, Melita Šalković-Petrišić

**Affiliations:** Department of Pharmacology, University of Zagreb School of Medicine, Zagreb, Croatia; Croatian Institute for Brain Research, University of Zagreb School of Medicine, Zagreb, Croatia; Health Centre Zagreb – West, Zagreb, Croatia; M3 Research Center, Excellence Cluster ‘Controlling Microbes to Fight Infections’ (CMFI), Interfaculty Institute of Microbiology & Infection Medicine Tübingen (IMIT), Eberhard Karls University & University Hospital Tübingen, Tübingen, Germany; German Federal Institute for Risk Assessment, German Centre for the Protection of Laboratory Animals and Experimental Toxicology, Berlin, Germany; University of Zagreb Faculty of Electrical Engineering and Computing, Zagreb, Croatia

**Author notes:** Corresponding author: Davor Virag, Department of Pharmacology University of Zagreb School of Medicine Zagreb, Croatia.

**Keywords:** Experimental Animal Models, Home Cage Monitoring, Bioethical Issues, Electronic Equipment and Supplies, Operant Conditioning, Open Source, Alzheimer Disease, Streptozotocin

## Abstract

Home cage monitoring (HCM) captures longitudinal animal behavioural data without human intervention. However, the systems’ complexity is rarely addressed in their design, increasing the risk of data loss, which wastes workhours, resources, and animal lives.

To assess the feasibility of implementing modern, robust architectures in complex operant HCM paradigms, the **V**ersati**L**e **A**utonomous **D**ev**I**ce for **S**cheduled **L**earning **A**ssessment **V**ia Wi-Fi (VLADISLAV) was developed and employed to test cognitive deficits in the intracerebroventricular streptozotocin-induced rat model of sporadic Alzheimer’s disease (sAD). Reliability was modelled against a system architecture common in commercial HCM systems by modelling the failure rate of the devices’ critical components across typical durations of animal experiments.

VLADISLAV assessed multiple cognitive dimensions of a rat model of sAD with automated, scheduled testing. Its design enabled simultaneous, redundant recording to multiple devices in real time, as well as batch remote control and supervision of tens of VLADISLAVs. VLADISLAV is estimated to reduce component failure rate ∼200-fold at €40/device.

Data loss due to system failure shouldn’t be accepted as a normal occurrence and robust system design is an ethical imperative. VLADISLAV’s robustness and utility demonstrate the potential of embedded networked systems, used in other industries and consumer electronics for over a decade. Today, the open source ecosystem enables cost-effective implementation of such architectures in HCM by biomedical researchers with no electronic engineering education, preventing data loss and facilitating researchers’ and technicians’ day-to-day work. Considering these findings, it is apparent that the implementation of modern architectures in HCM is long overdue.

## Introduction

Rodent behaviour testing is widely used in biomedical research in exploring disease pathophysiology and assessing novel therapeutic approaches. Technological advances and the widespread use of computers and microelectronics have refined the application of existing paradigms but also enabled the use of novel ones which previously weren’t feasible or even possible. One such change is increasing popularity and use of home cage monitoring (HCM) (Kahnau et al., 2023).

Automated monitoring in the home cage is a good complement, able to provide continuous, longitudinal data of tests performed in the home cage, minimising the confounding element of experimenter presence (Voikar & Gaburro, 2020). HCM can produce large, comprehensive animal behavioural datasets recorded in well-controlled, more replicable conditions. Amassing such datasets could improve translation through both analytic and exploratory approaches (Anello & Fleiss, 1995): i) in the former, facilitating preclinical meta-analyses can clarify animal phenotypes and the way the environment impacts them (Kafkafi et al., 2018; Mandillo et al., 2008); ii) in the latter, an -omics-like approach (including bioinformatic and “AI” techniques) to behavioural data can reveal previously unnoticed behavioural features of distinct pathophysiology (Bains et al., 2025; Baran et al., 2021; Kafkafi et al., 2018).

From an ethical standpoint, potential increase of reproducibility, reliability, and the amount of data extracted from preclinical studies directly aligns with the *Reduction* principle of the 3Rs of ethical animal research (Grieco et al., 2021; Kafkafi et al., 2018; Mandillo et al., 2008). Continuous welfare monitoring and alarms can prevent contamination by reducing the number of required physical entries into habitats, especially important in preventing potentially large losses of animals in germ-free and gnotobiotic facilities. This capability can also help identify subtle and early signs of distress (Fuochi et al., 2024), thereby contributing to both the *Reduction* and *Refinement* principles. Finally, more data could lead to better simulation of some facets of rodent behaviour, eventually contributing to *Replacement* (Fuochi et al., 2024; Rose et al., 2025; Zhang et al., 2025).

Notably, time budgeting follows different principles with automated HCM devices. An initial time investment in setup, proportional to the complexity of the electronic HCM system, frequently pays off through time saved via automation throughout the study (Kahnau et al., 2023). As the saved working hours can be allotted to other aspects of the experiment, it can be argued that less required labour could lead to more rigour while conducting experiments.

Alongside the additional layer of ethical responsibility, designing and conducting animal studies comes with a unique challenge. Beginning the *in vivo* stage of animal experiments is akin to flipping an hourglass – animals age, evolve pathophysiology in disease models, and retain previous experiences (e.g. behavioural tests, equipment malfunctions). Additionally, given the inherent variability of living organisms, unexpected events aren’t rare and may lead to inconclusive data. As “the hourglass” makes true technical replicates (analogous to repeating a molecular assay) virtually impossible, these events need to be addressed promptly and thoroughly during the study (Howard, 2002). Failing to attain the required data usually requires repeating the experiment, necessitating additional, considerable investment of funds and work hours, and, most importantly, the use of even more animals.

In HCM systems, the risk of unexpected events causing data loss is compounded by a greater chance of such events occurring in the systems’ continuous and unattended operation. As real-life conditions continuously change and frequently deviate from our expectations, it is difficult to predict myriads of ways in which an HCM system could fail.

Either way, these experiences may be a driving force for researchers to build custom HCM systems – either flexible enough to be rapidly adjustable in changing conditions, or optimised for a subset of specific circumstances. In addition or in place of these reasons, affordability can be an important factor due to high costs of most commercial systems (Habedank et al., 2022; Oellermann et al., 2022; Salem et al., 2024; Virag et al., 2025; Voikar & Gaburro, 2020). In a review of HCM-related publications by Kahnau et al., the formation of a larger market of HCM systems can be observed, beginning in the late 1990s. However, custom-built systems remain steadily used until the late 2010s when their use increases as well, coinciding and likely driven by a surge of publications utilising more user-friendly open electronics platforms released in the early 2010s (Kahnau et al., 2023; Oellermann et al., 2022).

Nowadays, many researchers with little to no formal education in the technical sciences successfully build HCM systems which yield useful and accurate data, thanks to the open source ecosystem (Habedank et al., 2022; Oellermann et al., 2022; Virag et al., 2025). Having access and insight into the inner workings of the device and without relying on a third party, researchers can rapidly refine and adapt the system through day-to-day use as both the operation of the system and what is required of it become clearer. In a process arguably similar to that described by Sharma et al., extensive theoretical expertise in electrical engineering likely isn’t crucial (Sharma et al., 2025). Indeed, publications of open source HCM systems generally emphasise practical advice relevant to everyday use, including limitations, relegating technical details to supplements and repositories (Brown et al., 2016; Habedank et al., 2022; Matikainen-Ankney et al., 2021; Salem et al., 2024; Singh et al., 2019; Taghipourbibalan & McCutcheon, 2026; Virag et al., 2025). Importantly, these publications include extensive validation data from real-world experiments obtained following the scientific method. Open source systems are transparent and affordable in material, but in cases of failure, the researchers’ only guaranteed resource is thorough information on the inner workings on the system, though other avenues exist. Assistance can be sought from the creators directly, and fellow users via platforms such as TheBehaviourForum.org. This avenue can be especially fruitful with mature, actively maintained, and widely used open source systems, though not many exist (De Chaumont et al., 2019; Matikainen-Ankney et al., 2021).

Commercial HCM systems are generally moderately to extremely expensive and nearly always closed source, but the purchase yields support by the company alongside the device itself. Therefore, solving challenges in everyday use is done via easy and direct access to personnel specifically trained to maintain and continuously develop the system in question. An investment in a pre-made system with support, if economically possible, can have positive ethical implications – if the researchers are relieved of the physical and mental work of designing, building, testing, and repairing a device, or negotiating outside expertise to do so, they can devote the immense saved time to other aspects of experimental design and execution, potentially improving them further.

The most prominent trend in consumer electronics over the past 15 years has been the widespread implementation of wireless connectivity, known as the Internet of Things (IoT) (Shehu Yalli et al., 2024). On the other hand, most commercial HCM systems are powered by personal computers (PCs) (COST TEATIME, 2025; COST-TEATIME, n.d.; European Cooperation in Science and Technology, 2025; Kahnau et al., 2023). PCs are not designed for continuous, unattended operation in environments such as animal habitats (De Micco et al., 2020; Koopman, n.d.; Matias et al., 2014) and such inappropriate use increases the risk of system failure. Replacing PCs with more appropriate electronics, especially small (“embedded”) networked electronics has shown to be exceptionally powerful in rare commercial systems (Kravitz, 2022; Slivicki et al., 2023; Tir et al., 2025a) and the open source modification of the widely used home cage operant conditioning Feeding Experimentation Device version 3 (FED3) (Matikainen-Ankney et al., 2021), the Realtime and Remote FED3 (RTFED) (Taghipourbibalan & McCutcheon, 2025).

The discussion of such architectures rarely centres on their immense potential for preventing data loss. Furthermore, there are practically no in depth discussions of HCM device robustness – a PubMed search for terms related to reliability (“*((“mean time to failure”) OR (“mean time between failures”) OR (mttf) OR (mtbf)) AND (“Animal Experimentation”[MeSH Terms])*”) yield no results.

Architectures which could increase robustness have been in widespread use for over 15 years but have hardly seen implementation in HCM. While the inertia may be caused by a lack of market pressure, it can be argued just as well that the research and development costs cannot be justified or sustained. Indeed, while a risk of data loss may be theoretically argued, is the magnitude truly concerning, or even practically relevant? More plainly put, are “obsolete” and “inappropriate” architectures in HCM system designs simply good enough?

Due to the ethical implications of data loss in animal studies, this study aims to open an evidence-based discussion of robustness in HCM devices through a case study of designing, building, and testing a novel device with the discussed principles in mind.

In this study, the *VersatiLe Autonomous DevIce for Scheduled Learning Assessment Via Wi-Fi* (VLADISLAV), a networked home cage operant conditioning device was built with data loss prevention as the primary design objective. Secondary objectives were practicality and low cost. VLADISLAV was then employed in a proof-of-concept study of a rat model of sporadic Alzheimer’s disease to demonstrate its effectiveness and evaluate its practicality. VLADISLAV’s robustness and practicality were quantified and compared to a common commercial system for operant conditioning in the home cage. Finally, the findings are discussed to establish whether the excess risk of currently widespread HCM architectures is of practical significance, and whether it justifies the cost and effort of a push towards more robust HCM architectures.

## Materials and Methods

### Building VLADISLAV: A Simple Embedded Networked System for Operant Conditioning in the Home Cage

VLADISLAV is an automated, remotely operated home cage operant conditioning (HCOC) device which attaches to standard rodent cages. It performs unattended, rodent-initiated testing based on a flexible protocol, determined by the experimenter’s specific research question. The protocol definition and trial results are exchanged remotely via Wi-Fi.

#### Apparatus

The VLADISLAV apparatus consists of three main components:

##### The nosepoke opening (Figure 1 and Figure 2; Label A)

A square aluminium profile contains a beam-break sensor which forms an infrared beam between an LED transmitter and a receiver – when this beam is obstructed by a rodent nose, the microcontroller registers a nosepoke event. The microcontroller board is mounted behind the nosepoke opening, with its onboard LED providing light cues through the nosepoke opening.

**Figure 1.**
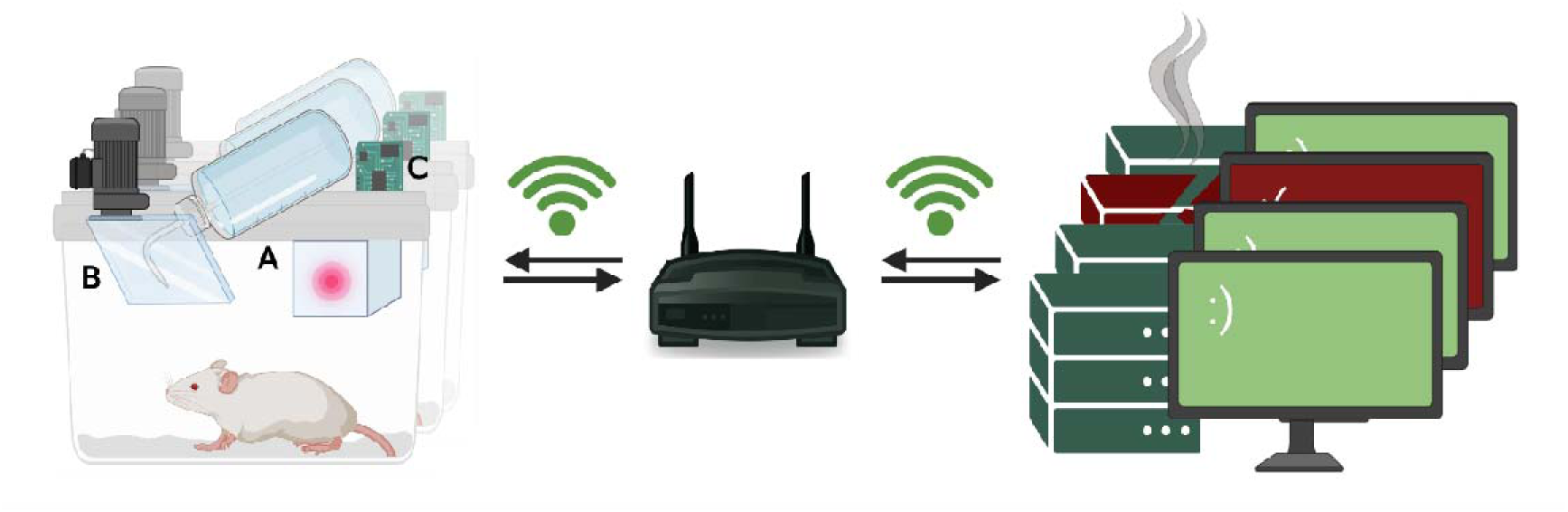
A schematic of VLADISLAV’s main components and network architecture. *Note.* A. An illuminated nosepoke opening, which the rodent uses to operate the device according to its light cues. B. A motor-operated bottle flap, gating access to drinking water as a reward. C. An Arduino Wi-Fi-enabled microcontroller board, which conducts the test and transmits results wirelessly.

**Figure 2.**
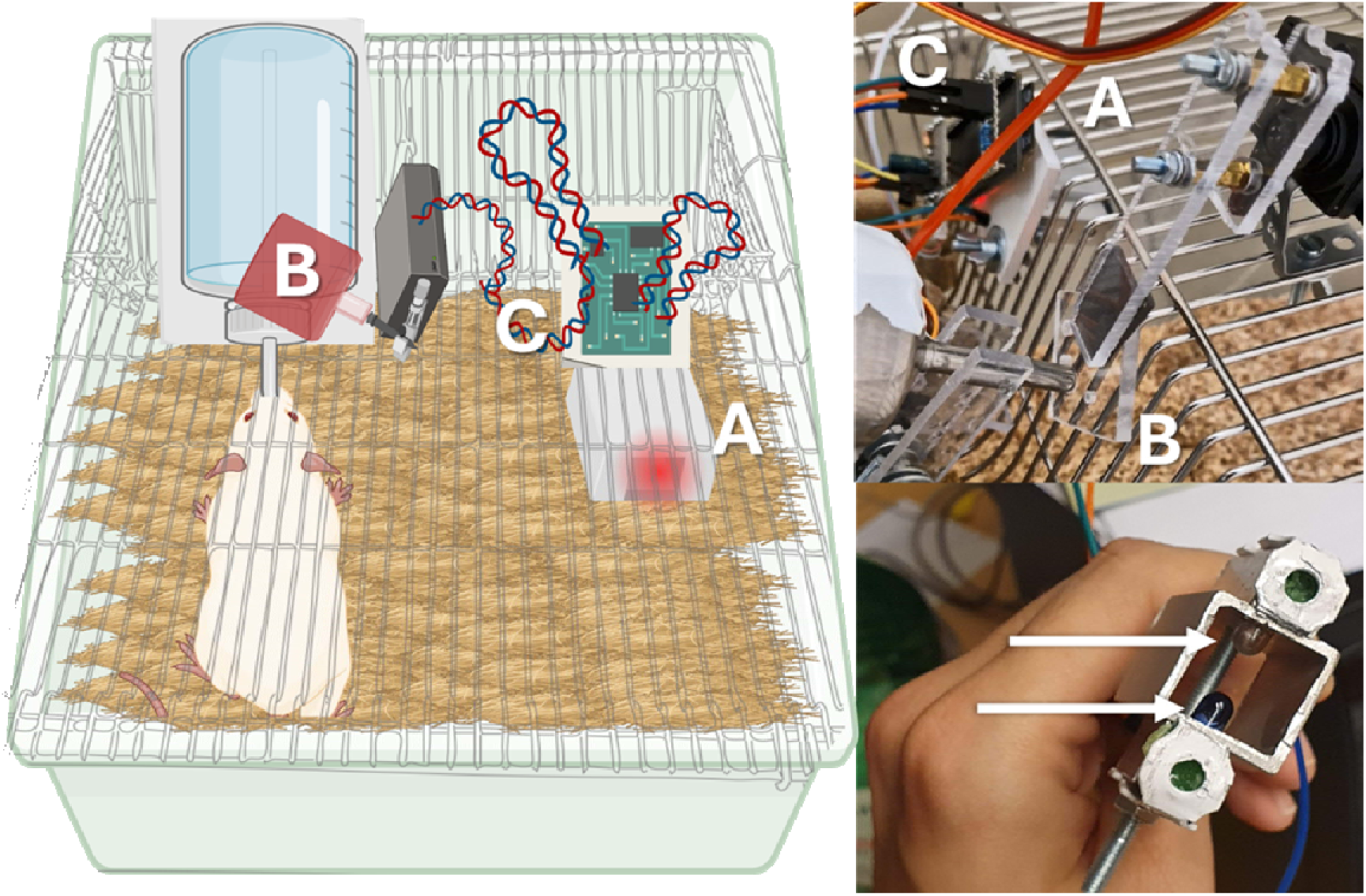
The VLADISLAV testing apparatus. Note. The left panel shows a top view of the positioning of the apparatus’ main components: the nosepoke opening within the cage (A), the servo motor-operated water gate (B), and the microcontroller board with an onboard LED glowing red (C), illuminating the nosepoke opening (letters marked in white). In the top right panel, the same components are shown as part of a fully assembled, operational VLADISLAV. As part of its operation, the microcontroller (C), in reaction to an appropriate nosepoke response (A), engages the servo which opens a gate (B) giving access to the water/reward. In the bottom right panel, a close-up of the nosepoke opening is shown from the cage aspect. The white arrows mark an infrared transmitter and receiver pair which forms the beambreak sensor used to detect nosepokes.

##### The motor-operated bottle flap (Figure 1 and Figure 2; Label B)

Rewards are dispensed by granting access to drinking water. A simple bottle holder keeps the tip of the spout just outside of the cage lid, where it is accessible to the rodent, but can be obstructed by a small plastic panel. The movement of the panel is performed by an MG995 servo motor.

##### The microcontroller and the network architecture (Figure 1 and Figure 2; Label C)

The microcontroller setup is based on the *Multicage InfraRed Open-Source Locomotor Activity eValuator* (MIROSLAV) home cage platform (Virag et al., 2025) – a Wi-Fi-enabled Arduino-compatible microcontroller board, ESP32-S2-Saola-1, continuously sends status and trial updates and can receive and permanently store new protocols at any time wirelessly, via the Message Queuing Telemetry Transport (MQTT) communication protocol (Banks et al., 2019). This enables logging and control with a theoretically limitless number of on- and off-site network-connected computers. A computer is not required in the habitat, and Wi-Fi coverage is sufficient for complete VLADISLAV functionality. Wi-Fi is also used for precise time synchronisation against a Network Time Protocol (NTP) server, allowing the testing protocol to be initiated automatically at a precise time of day, or day of the week.

##### Assembly

VLADISLAV’s architecture utilises a simple design with few components to facilitate novel assemblies appropriate for each laboratory’s use cases. The assembly used in our studies is shown in Figure 2, with all parts internal to the cage made from aluminium as we have found that, generally, plastic materials cannot withstand the rats’ interest over the course of one night. More details on the assembly can be found in Supplementary material 2 and in the VLADISLAV repository (Virag, 2026). The total cost of 40 cage devices is around €1600 (including power supplies and Wi-Fi routers).

##### Power supply

As the full experiment involved MIROSLAV monitoring, each cage was equipped with a powered MIROSLAV cage unit (Virag et al., 2025). The cage units were amended with a pin header to break out the power connections, which were then used to directly power each cage’s VLADISLAV via jumper wires. Without concurrent use with MIROSLAV, a larger number of VLADISLAVs may be conveniently powered using standalone MIROSLAV power distribution boards (Virag, 2024) with appropriate connectors. A VLADISLAV can also be powered via the onboard USB connector, particularly useful with a smaller number of VLADISLAV devices or while prototyping, as the connector is also used for programming access.

#### Protocol

##### Two-signal operant conditioning

Fundamentally, the protocol is an operant conditioning paradigm similar to the Variable Delay-to-Signal (VDS) test (Leite-Almeida et al., 2013). When the apparatus initiates a testing session, the water gate is closed, and the LED illuminates the nosepoke opening with a blue colour. The rodent initiates a trial by responding to the first signal, the “pre-cue”, by performing a nosepoke. The illumination ceases, and after a delay (e.g. 3-12 seconds), the “cue” follows for a specified duration (e.g. 5 seconds) as a blinking yellow light. If the rodent performs a nosepoke, the light turns off, the water gate opens and the trial is counted as a success, and a failure otherwise. Finally, a new trial begins with a blue illumination of the nosepoke opening, and this cycle repeats until the end of the testing session. At the end of each session, illumination is stopped, and the water gate opens.

The test stage is preceded by two training stages, during which the rodent learns to operate the nosepoke to gain access to water, and to respond to the nosepoke light cue, respectively. In these sessions, trials are initiated automatically, and only the “cue” signal is used, with one nosepoke being sufficient to grant water access.

Per day, one or more sessions are performed, typically 1-4 hours in duration depending on session type (training or test). After the scheduled daily protocol is complete, the device provides unrestricted water access.

##### Protocol control

The testing schedule and protocol parameters can be set at any time remotely, via Wi-Fi, and the current protocol can be examined using a web browser. Pre-set templates are also supported, which enable the experimenter to load an entire predefined protocol (e.g. testing/training, easier/more difficult) with a single command. A session can be stopped and reinitiated by remote command, loading newly-set values. To support scalability in larger or multiple simultaneous experiments, devices may be controlled and managed as a group. Any command sent to a group is distributed and simultaneously executed by all devices assigned to it. Devices emit status and experimental event messages (e.g. nosepokes) immediately as they occur, and results of each trial and session as they are completed, enabling real-time monitoring of the rodents’ performance.

Other paradigms may also be implemented by modifying the Arduino code, available in the VLADISLAV repository (Virag, 2026).

#### Data preparation and validation

Data validity is ensured with the *Prepare-a-VLAD* Python notebook, available in the VLADISLAV repository (Virag, 2026). Additionally, the notebook can be used to add metadata and perform exploratory analyses with grouping, filtering, and heatmap visualisation.

Further information may be found in Supplementary material 2.

### A Proof-of-Concept Study: Cognitive Deficits in a Rat Model of Sporadic Alzheimer’s Disease

VLADISLAV was used as a part of an experiment aimed to assess the effects of a novel treatment approach in the streptozotocin-induced rat model of sporadic Alzheimer’s disease (AD) (manuscript in preparation). In this work, a subset of the data is presented, which focuses on the cognitive performance of streptozotocin-treated animals in comparison with the controls in VLADISLAV testing.

#### The Rat Model of Sporadic Alzheimer’s Disease

The model of sporadic AD used in this study was initially introduced by Mayer et al. and Lacković & Šalković (Lacković & Šalković, 1990; Mayer et al., 1990; Salkovic-Petrisic & Hoyer, 2007). Streptozotocin (STZ), a diabetogenic compound, is injected into the lateral ventricles (intracerebroventricularly – icv) of the rat brain following a standard procedure (coordinates –1.5 mm posterior, ±1.5 mm lateral, and +4 mm ventral to the bregma (Noble et al., 1967)), subsequently optimised by our group (Homolak et al., 2021, 2023; Salkovic-Petrisic et al., 2021). Briefly, STZ (1.5 mg/kg) is freshly dissolved in 0.05 M citrate buffer (pH 4.5) prior to application, and a volume of 2 μL is injected into each ventricle, with a final dose of 3 mg/kg (Homolak et al., 2023; Knezovic et al., 2017).

Twenty male Wistar rats 3 months of age were assigned to two treatment groups based on stratified randomisation with regard to body weight, litter, and home cage placement (described below). The experimenters were not blinded, with the limitation’s rationale described in Supplementary material 1. Ten animals were injected with the STZ solution (N_STZ_ = 10), while, by the same procedure, another ten were injected with citrate buffer alone to serve as the control group (N_CTR_ = 10) under ketamine/xylazine anaesthesia (70/7 mg/kg) administered intraperitoneally. The procedure was carried out on both days from ZT (Zeitgeber time) 2 to ZT 4.

In the literature, STZ-icv-treated rats eventually develop cognitive decline, the onset of which can be detected around 2 weeks post treatment (Salkovic-Petrisic et al., 2022).

#### Animal Housing

Animals were bred and kept at the licensed facilities of the University of Zagreb School of Medicine’s Department of Pharmacology (HR-POK-007) and Croatian Institute for Brain Research (HR-POK-006), respectively. The facilities were kept at 21-23°C and 40-70% relative humidity. The 12-hour light cycle began at 06:00 (ZT 0) and ended at 18:00 (ZT 12). At the age of 3 months, the animals were single-housed, starting 7 days prior to the STZ treatment. Single housing was performed due to the requirements of the full experiment outside the scope of this work: administration of treatment in drinking water to additional groups and individual VLADISLAV testing. The cages of each treatment group were placed on two racks top-to-bottom by body weight (1^st^ rack: animals 1-5; 2^nd^ rack: animals 6-10). The cages were changed with fresh bedding (Scobis Uno, Due; Mucedola S.R.L., Italy) once per week and no enrichment or nesting material was used.

The animals were fed with standard pellets (Mucedola S.R.L., Italy) and food access was unrestricted. Starting 10 days after completing the STZ model induction, the animals were mildly water-deprived, with water access withheld during the light phase from 10:00 to 18:00 (ZT 4 to ZT 12), conditioned on VLADISLAV task-solving from 19:00 to 23:00 (ZT 13 to ZT 17). Due to a special treatment protocol applied in the full experiment, but outside the scope of this work (manuscript in preparation), the animals’ bottles were filled with 33±5 mL of water (adjusted for body weight, with overhead for spillage) at 18:00 (ZT 12). The remaining water was noted for each animal at 7:00 (ZT 1) to adjust the volume for the subsequent night, with the goal of a few millilitres left over, signifying that the volume of water was sufficient throughout the night, but nearly completely consumed to ensure delivery of dissolved treatment in groups not described in this work. Immediately thereafter, free access to water was given from 7:00 (ZT 1) to 10:00 (ZT 4).

Visual inspections were carried out by licensed personnel twice per day at 7:00 (ZT 1) and 18:00 (ZT 12).

#### VLADISLAV Home Cage Testing

VLADISLAV testing began one week after STZ model induction. The devices were activated at the beginning of each dark period (19:00, ZT 13) every day for 24 days. The protocol was modelled after the VDS paradigm (Leite-Almeida et al., 2013), spanning three consecutive phases as shown in Figure 3: easy training, difficult training, and the test. Devices were active each night for 6 hours during the training phase, and 4 hours during the testing phase.

**Figure 3.**
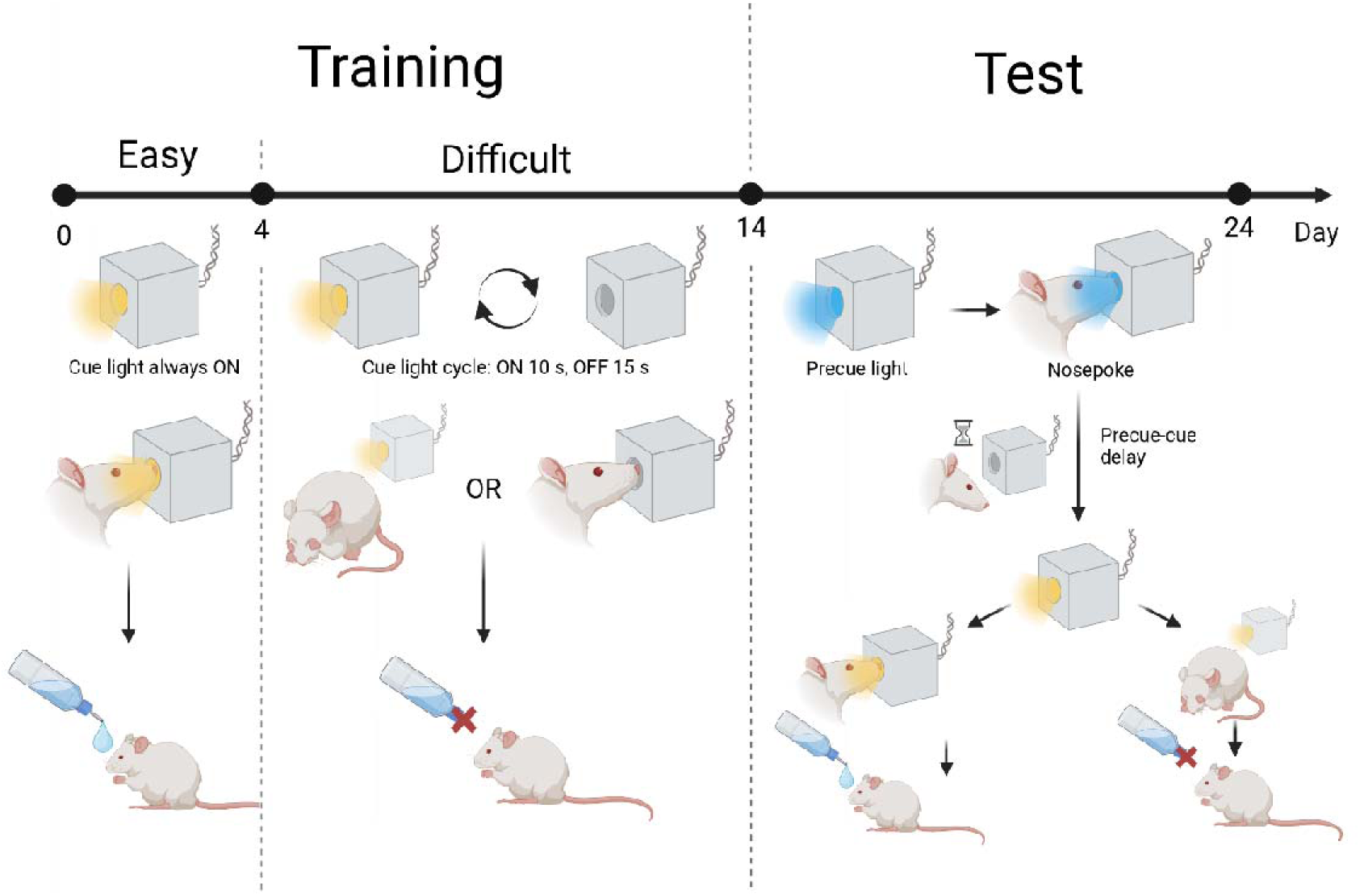
Protocol of VLADISLAV home cage testing. Note. Training, easy (day 0-4): The flashing cue light, shown in yellow, was constantly on throughout the testing period, and a single nosepoke yielded the reward. Training, difficult (day 5-14): The flashing cue light was on for 10 seconds, and off for 15 seconds. Only nosepokes to the cue light were rewarded. Nosepokes while the cue light was off did not yield a reward, and would reset the 15-second off period, delaying the following cue presentation. Test (day 15-24): The precue light, shown in blue, was on until the animal responded with a nosepoke, or until the daily testing period elapsed. A response initiated a trial and resulted in cue presentation after the precue-cue delay elapsed. The flashing cue light was presented for 40 seconds, and an additional, second nosepoke was required during this period to obtain the reward.

During the first four days, the flashing cue light (pictured as red illumination in Figure 3) was continuously on, with every nosepoke granting a reward (training, easy) to establish a relationship between nosepoking and a reward (access to water).

From day 5 to day 14, the OFF period for the cue light was introduced (training, difficult). The flashing cue light was on for 10 seconds and off for 15 seconds, with this sequence repeating in a loop. While the cue light was off, nosepokes did not yield a reward to more specifically associate the nosepoke-obtained reward with the flashing cue light.

On day 15, when the majority of the animals crossed the criterion of 85% correct responses, the test phase began, which required the animal to manually initiate a trial as opposed to automated continuous cycling. While the criteria for receiving a reward remained the same as in the difficult training phase, the precue light was on until the animal responded with a nosepoke, or until the daily testing period elapsed. A nosepoke would start the variable precue-cue interval (light off), followed by a trial as in the difficult training, starting with the flashing cue light. Nosepokes during this precue-cue interval were counted as “impulsive nosepokes” and did not yield a reward. Moreover, these nosepokes would reset the precue-cue interval, thereby delaying the start of a trial (i.e. appearance of the flashing cue light). If there were no interactions during precue-cue interval, the flashing cue light was then presented for 40 seconds, and a response to the cue yielded 15 seconds of access to drinking water as reward. After the reward period, the precue light would again be presented, with VLADISLAV ready for the rodent to initiate a new trial.

Each night during the test protocol (days 15-24) was split into three sessions, differing in the precue-cue delay. The first and third session spanned the first and last fifth (i.e. 48 minutes) of each night (4 hours from 19:00 to 23:00, ZT 13 to 17) and were configured to use a precue-cue delay of 3 seconds. The second session, corresponding to the middle three fifths (144 minutes) of the night, used precue-cue delays of either 6 or 12 seconds, chosen randomly for each trial and noted in the logs.

The goal of the training stages was to train the animals to operate the nosepoke opening in the context of the cue light. The testing protocols introduced a significant change in the task, where animals were required to start trials on their own, and then needed to withhold nosepoking during the precue-cue delay. In the second session of each night, the cue is presented after a longer, variable delay, which presents a further complication in the learning process, and is designed to elicit impulsive responses as per the VDS paradigm (Leite-Almeida et al., 2013).

#### Other In Vivo Procedures and Sacrifice

During the experimental course, a sequence of standard out-of-cage tests was repeated three times in two-week intervals to assess different facets of behaviour: i) open field (Seibenhener & Wooten, 2015), ii) novel object and location recognition (Antunes & Biala, 2012), and iii) social interaction (Jabarin et al., 2022). Before sacrifice, the passive avoidance test (Ögren & Stiedl, 2013) was performed. The Prepare-a-VLAD notebook labels the dataset with the behavioural test performed on each day (if any) as part of the data preparation process.

Ultimately, the animals were sacrificed 7 weeks after STZ-icv treatment under deep anaesthesia (thiopental-diazepam 70+7 mg/kg) to obtain tissue samples as part of the full experiment outside of the scope of this work.

#### Data Cleaning and Statistical Analysis

Faults were detected using experimental notebook logs and VLADISLAV self-reports. The faults were a result of assembly errors or the animals’ interactions with the devices (e.g. forcing the gate open or chewing through improperly secured wires), with no failures attributable to device crashes or other aspects of the embedded system architecture. Data was cleaned to remove faulty devices or those with no animal interactions using the Prepare-a-VLAD notebook. Ultimately, data from 7 control animals, and 6 STZ-treated animals were analysed.

Statistical analysis was performed in R (R Core Team, 2024) with the *glmmTMB* R package (Brooks et al., 2017). Logistic regression was performed to assess test performance over time. A generalised linear mixed model with the binomial distribution was fitted to the binary (correct-incorrect) data. A three-way interaction was specified between polynomial time, treatment group, and session number as fixed effects, with an appropriate random effects structure. The three-way interaction allows the assessment of treatment effect both on overall performance, and performance across the three daily sessions over the experiment’s timespan. Probabilities are back-transformed from the logit scale. Confidence intervals were adjusted for multiple comparisons using the Šidák correction (Šidák, 1967) for the overall curves (Figure 4). Curves by session (Figure 5) may only be interpreted to generate hypotheses for future studies to avoid “Hypothesizing After the Results are Known” (HARKing) (Kerr, 1998).

**Figure 4.**
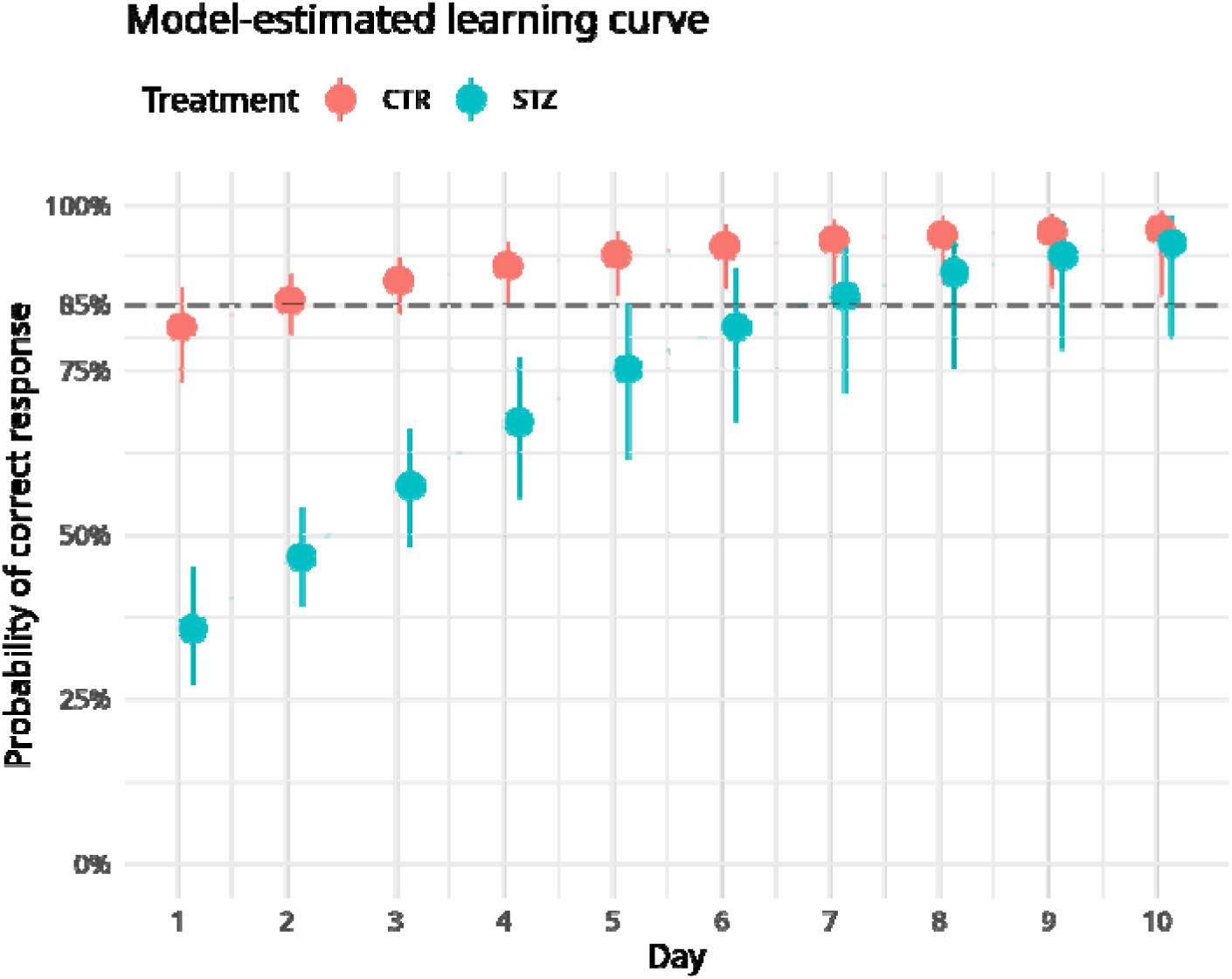
Performance on the test by treatment group over 10 days. Note. Learning curves of the control (red) and STZ-treated (blue) groups cross the 85% threshold on the second, and the seventh day, respectively. The points represent estimated marginal means, and the vertical lines 95% confidence intervals. (N_CTR_ = 7, N_STZ_ = 6)

**Figure 5.**
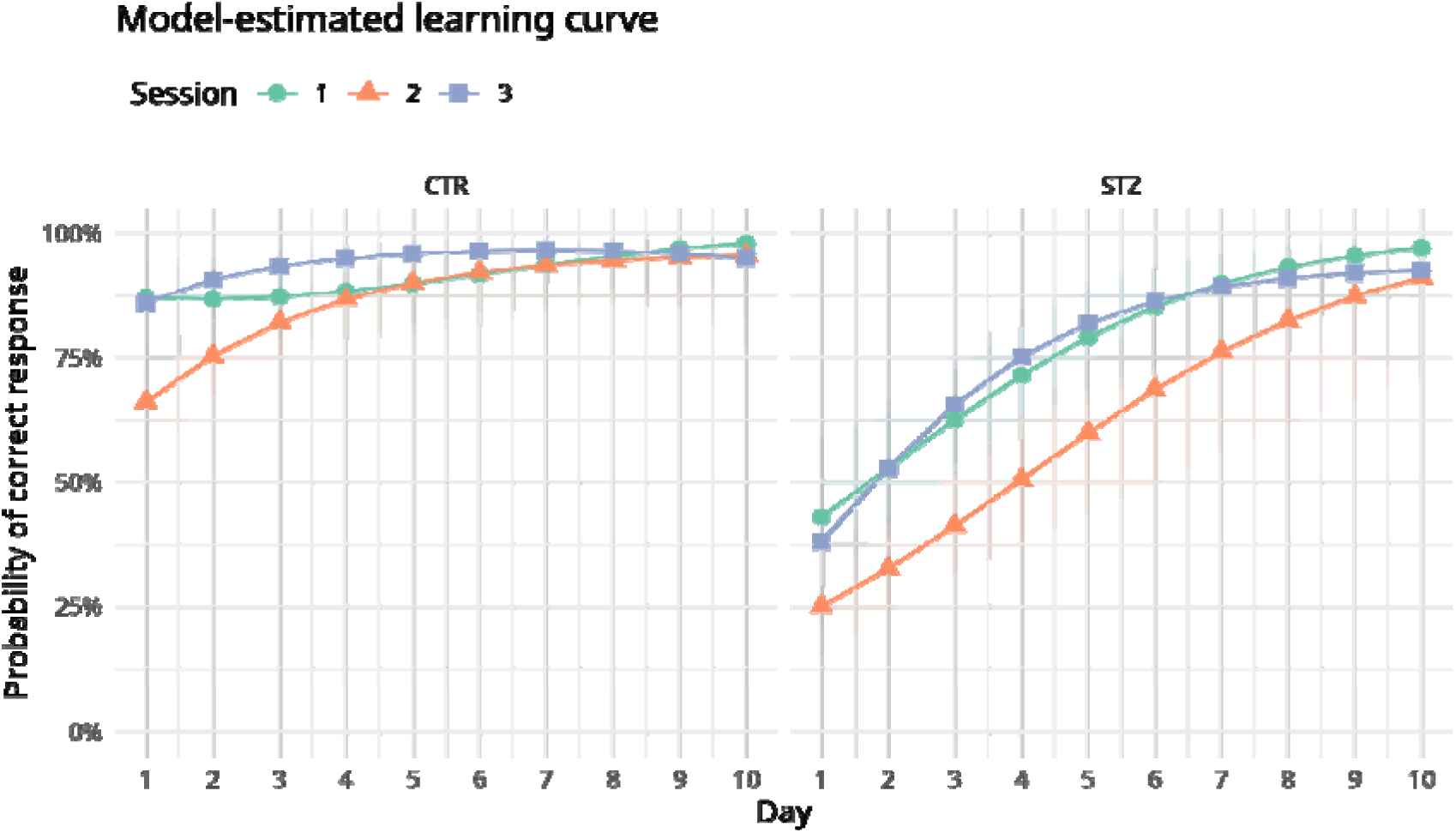
Performance on the Test by Treatment Group Over 10 Days, by Session. Note. The data from Figure 4 is stratified by session. Each night, the three sessions are performed in order, with the middle session (orange) applying a more difficult protocol than the first (green) and last sessions (blue) of the night. The STZ-treated group consistently tends to perform worse in the middle session until the final days of the protocol, while the control group’s performance in all sessions equalises within 3 days. The points represent estimated marginal means, and the coloured ribbons 95% confidence intervals. (N_CTR_ = 7, N_STZ_ = 6)

Though each individual trial’s outcome is passed to the previous model, another model was fitted with data binned per night for each animal to assess the number of initiated trials each night with regard to conventional behaviour testing performed during the day. A generalised linear mixed model with the negative binomial distribution was fitted to the daily trial count data, with interactions between treatment and polynomial time, and between treatment and a binary variable indicating whether conventional behavioural testing was performed prior to VLADISLAV testing on that day.

Model selection and diagnostics are performed based on visual inspection of fitted data and residuals using the *DHARMa* (Hartig, 2022) and *performance* (Lüdecke et al., 2021) R packages. Marginal means and contrasts are then calculated using the *emmeans* package. All plots are produced using *ggplot2* (Wickham, 2016).

### Estimating Reliability in VLADISLAV and a Common Commercial Apparatus

#### Impact of System Architecture on Reliability

Commonly, in HCM systems, the cage electronics send data to a PC via USB, which the PC saves to its local hard drive. The intended use usually implies that the researcher makes regular copies (backups) of the data to another device. The PC can fail in myriads of ways, with hard drive failure having the most extreme consequences. Upon failure, the PC will stop operating the cage electronics, and loss of all previously recorded data is very likely.

VLADISLAV cage electronics contain networking functionality and directly transmit data to a local Wi-Fi network powered by a router, which is then simultaneously distributed to a number of recording devices (PCs, servers, network-attached storage etc.). Thus, the cage electronics are autonomous, and data storage depends on the Wi-Fi router distributing data to PCs and other complex devices. Thus, the main design change is that the system only depends on so called “embedded” electronics (cage electronics, Wi-Fi router) – failures of any single computer involved will not stop device operation nor result in loss of data.

Though routers and PCs are electronic devices designed for different use cases, with routers being more appropriate for unattended, continuous operation in harsh environments, it is prudent to assess this assumption by comparing the reliability of a router and a PC, the central components of the two architectures. Directly testing device reliability is a complex electronic engineering task requiring specialised methodology (Gandhi et al., 2019; Martin, 1999; Whitman, 2003), however, a rough comparison can be made using sparse existing data in the literature.

#### Reliability Curves

To compare the risk of failure between a PC and a Wi-Fi router, two parameters were of interest: i) reliability, which corresponds to the probability of a device still operating without failure at a given timepoint, and is directly plotted over time as the reliability curve, and ii) mean time to failure (MTTF), which corresponds to the reliability curve’s area under the curve (AUC). MTTF may then be directly compared as a general, rough estimate of reliability of the two device classes, whereas the mathematical model of a reliability curve can provide data of the probability of a PC or a router still being operational at given timepoints.

An empirical reliability curve for consumer PCs, as those commonly found in HCM systems, was plotted from real life reliability data in the work by Matias et al. (Matias et al., 2013). All failures recorded in the study against which reliability was estimated have the potential to cause data loss, either as full system crashes, or system errors which can disrupt the device application. As the full dataset is inaccessible, datapoints of the curve were extracted manually from Figure 3 of the paper by Matias et al. (Matias et al., 2013) for PCs in the academic setting (“G1”), and a model was fitted using the *glmmTMB* R package (Brooks et al., 2017). Our modelled curve’s AUC (419.79 hours) roughly corresponds to the authors’ calculated MTTF (411.48 hours) listed in Table 8 of the source paper (Matias et al., 2013).

Reliability curves of common, commercial routers could not be found, likely due to this data being mostly collected in private companies’ internal stress testing laboratories and therefore proprietary. However, the MTTF is estimated at about 100,000 hours for routers – this figure may be lower for newer models which may have as-yet undiscovered design faults, and higher for older models in which these faults were fixed through experience from extensive field use (Kogan, 2010). The MTTF parameter corresponds to the AUC, which allowed us to rescale the reliability curve of PCs measured by Matias et al. (Matias et al., 2013) from an AUC of 419.79 hours to 100,000 hours, under the assumption that the actual reliability curve of routers is generally similar in shape to that of PCs.

#### Comparing Reliability Between PCs and Routers

Once mathematical models for both device classes’ reliability were obtained, the curves were trimmed and rescaled to start at 100% reliability at 1 hour elapsed, as it is expected that the systems will be heavily monitored and promptly repaired by the researcher at least within the first hour of use.

For practical relevance to basic and preclinical animal studies, the two models were used to calculate reliability at 1 months, 3 months, and 1 year. We also calculated the timepoints at which the reliability of the drops to 50% and at which the reliability of the router drops to 95%, followed by the reliability of the other device class (router and PC, respectively) at these timepoints.

## Results

### Proof-of-Concept Study

On the 21st day after the second STZ-icv application, the testing protocol was initiated for the subsequent 10 days. Figure 4 shows the binomial model-estimated mean probability of solving a task trial correctly. On the first night, the mean probability was 82% (CI_95%_ 73% - 88%) for the control group, and 36% for the STZ-treated group (CI_95%_ 27% - 45%) (OR_STZ/CTR_ 0.13; CI_95%_ 0.05 - 0.30). The control group’s mean performance surpassed 85% on the second day while it took a longer period for the STZ-treated animals that reached the 85% threshold on the seventh day.

In Figure 5, three learning curves are shown, one corresponding to the model-estimated correct response probability in each of the three daily sessions. During the first five days when the learning process is ongoing for both groups, the control group tends to improve over the duration of the night, performing better in Session 3 than Session 1 of the same night (CTR, day 4: OR_S3-S1_ 2.56; CI_95%_ 1.02 - 6.46), unlike the STZ-treated group (STZ, day 4: OR_S3-S1_ 1.21; CI_95%_ 0.70 - 2.09). In the control group, performance in Session 2 reaches that of Sessions 1 and 3 within three to four days, whereas this only occurs after more than one week in the STZ-treated group.

Overall, the STZ-treated group performs twice as many trials as the control group (OR_STZ/CTR_ 2.14, CI_95%_ 1.48 - 3.09). The number of trials is increased in the STZ-treated group from day 3 onwards (Figure 6A), and the difference lessens towards the end of the testing period when both groups reach a plateau in the learning curve. Furthermore, the number of trials is increased in both groups on days when conventional behavioural testing was performed prior to that day’s VLADISLAV test (Figure 6B), and this increase is far more prominent in the control group (Ratio [Behaviour 1 to 0 in STZ] to [Behaviour 1 to 0 in CTR] = -0.35, CI_95%_ -0.67 - -0.02).

**Figure 6.**
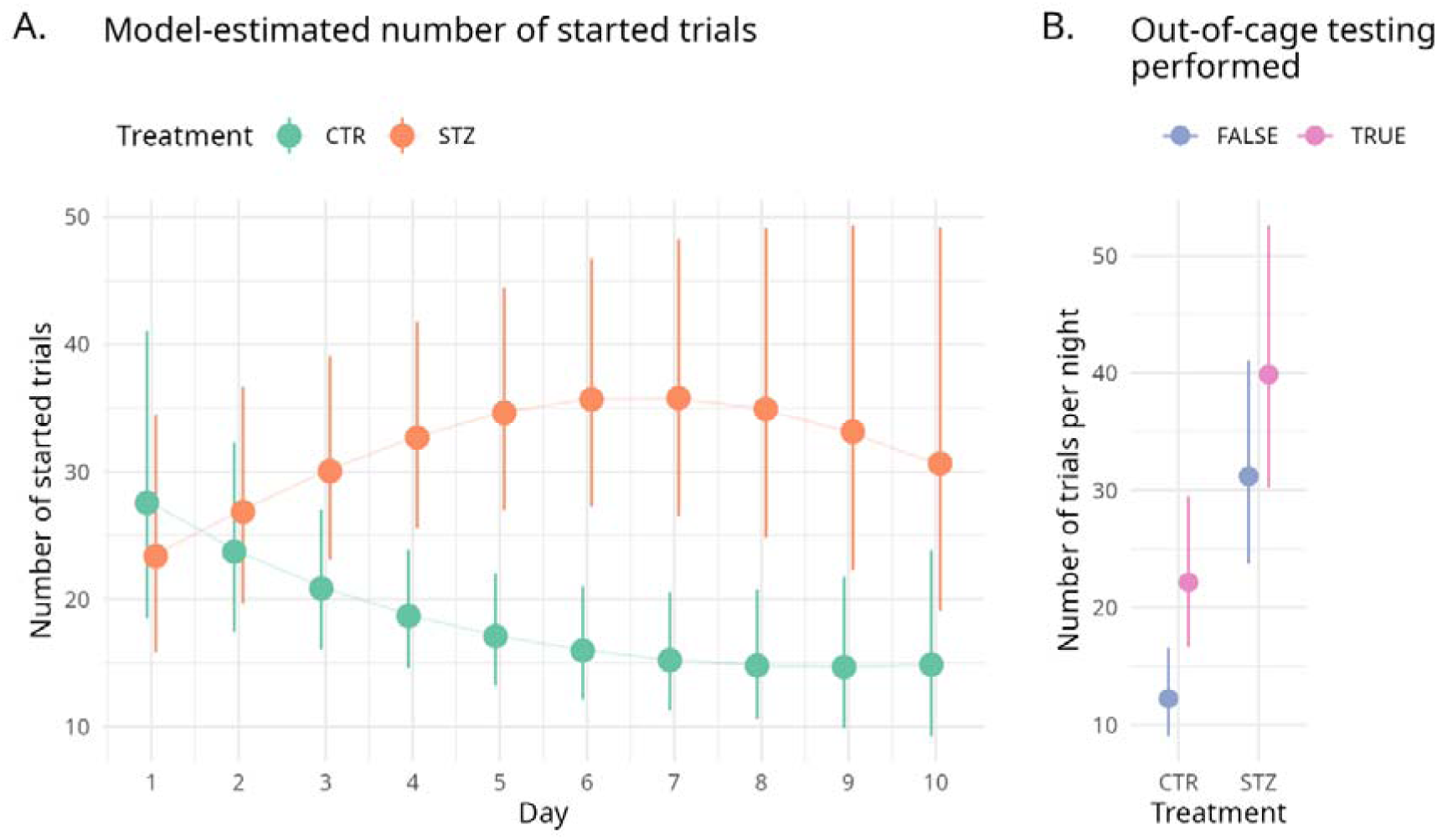
Number of Trials by Treatment Group. Note. In panel A, a time series plot is shown across the 10 days of the test. After the third day, the STZ group consistently performs more trials each night. Panel B shows the overall daily number of trials by group, depending on whether conventional, out-of-cage behavioural testing was performed during the day prior to VLADISLAV testing. The number of trials is increased, though to a lesser extent in the STZ-treated group. In both panels, the points represent estimated marginal means, and the vertical lines 95% confidence intervals in both.

### Comparison of Reliability Between VLADISLAV and a Standard Commercial Solution

Figure 7 shows modelled reliability curves for the PC and the router as single points of failure in a standard home cage operant system and VLADISLAV, respectively.

**Figure 7.**
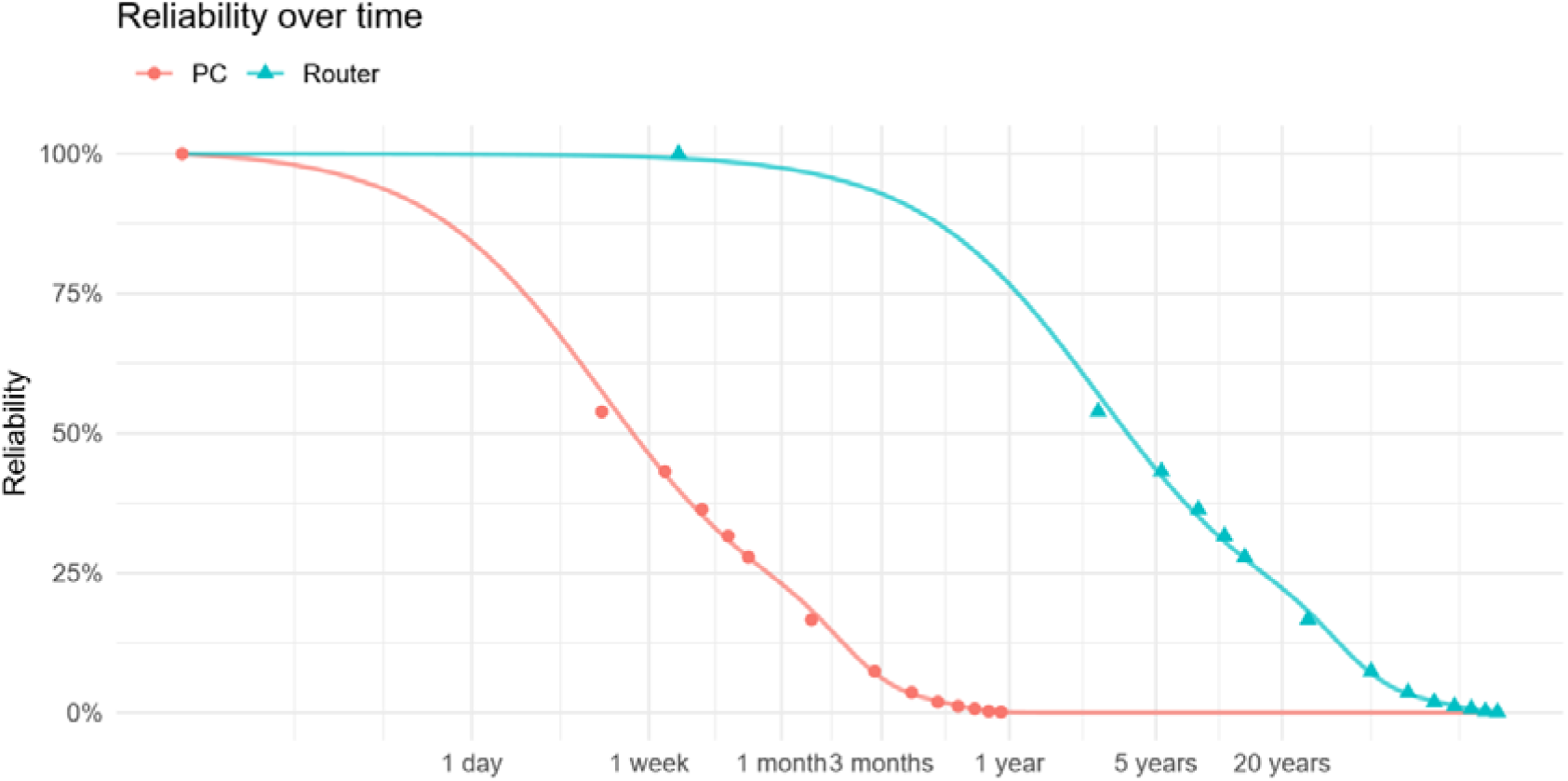
Modelled Reliability Curves for a PC and a Router as Single Points of Failure. Note. The points mark empirical reliability data extracted from the paper by Matias et al. (Matias et al., 2013) for PCs, and appropriately scaled for routers, as described in Materials and Methods. The lines represent predicted values from the best-fitting generalised linear model.

In Table 1, specific reliability values are extracted in the context of common experiment durations (1 month, 3 months, 1 year), as well as 50% PC reliability and 95% router reliability. After 3 months, it is expected that over 90% of PCs will have experienced a failure, whereas over 90% of routers will have been in continuous operation. The limitations of this methodology are further discussed in Supplementary material 2.

**Table 1.** Modelled Reliability Values at Specific Timepoints.

| Time point | Reliability |  |
| --- | --- | --- |
|  | PC<br>(used in standard device) | Router<br>(used in VLADISLAV) |
| 1 month | 23.06% | 97.46% |
| 3 months | 6.16% | 92.80% |
| 1 year | 0.07% | 76.88% |
| 5.87 days | 50.00% | 99.49% |
| 2.03 months | 11.88% | 95.00% |

*Note.* Reliability of both devices was estimated at timepoints of practical relevance for animal experiment durations utilising home cage systems (1 month, 3 months, 1 year). Additionally, the timepoints of 50% PC reliability and 95% router reliability were calculated, with reliabilities of the opposing device class at those timepoints.

## Discussion

### Experimental Findings with VLADISLAV

VLADISLAV has provided insight into a severe learning deficit of the rat STZ-icv model of sporadic AD in an operant conditioning task, which, to our knowledge, is the first such characterisation in the home cage.

Furthermore, the STZ-treated group performs about twice as many trials as the controls overall. When assessing the number of trials in the context of the learning curve, it is possible that animals perform more trials until they reach a plateau in learning. The plateau occurs within the first few days in the control group, and within the last few days in the STZ-treated group, and a fall in the number of trials is observed within each group during these periods, respectively.

In line with previous findings in the literature (Bruinsma et al., 2019; Kahnau et al., 2022), this study corroborates, and generates additional evidence that dramatically shorter training times (i.e. days) can be obtained in the home cage variant of an operant conditioning task. Furthermore, only drinking water was used as a reward, facilitated by <12 hours of water deprivation in the light phase. Observations regarding the necessity of water deprivation are detailed in Supplementary material 1.

Findings of this study should be interpreted in the context of its limitations – with a lack of cage enrichment and in social isolation (carried out to avoid possible impact on assessment of cognitive abilities and to ensure adequate dosing of the tested compound), the rats are likely understimulated, which could impact interaction with VLADISLAV. More information about the study design, including further discussion of the study’s limitations unrelated to VLADISLAV is available in Supplementary material 1.

### Managing Tens of Home Cage Devices Simultaneously

To effectively deploy a home cage operant device in the ongoing research project, the device(s) needed to be usable for unattended, scheduled testing of up to 40 single-housed animals (Note: Single housing as a limitation is discussed in Supplementary material 1). For a device to be considered “usable”, it needed to be designed with an architecture that could easily sustain a larger deployment, allowing easy simultaneous management of tens of devices while minimising the risk of data loss. Thus, VLADISLAV was developed based on our previous open source platform, MIROSLAV (Virag et al., 2025), a locomotor activity monitoring device with networking implemented to increase robustness. Reimplementing MIROSLAV in the context of operant conditioning as VLADISLAV has shown additional, remarkable utility.

The networked architecture provided immediate, real-time access to experimental and status logs at any time without entering the habitat. Since the proof-of-concept study was also part of a larger project, also testing simultaneous deployment of 40 VLADISLAVs for the first time, many failures were caught, as is expected while refining a prototype during a first-time large scale deployment. As previously mentioned, zero failures were related to the networked architecture showcased in VLADISLAV or its components. In the experiment, networking allowed quick detection, diagnostics, and resolution of failures, with most of the process done remotely, minimising disruption of the animals.

To define operant task protocols more easily, VLADISLAVs were programmed with protocol templates. An entire protocol can be loaded, or specific parameters (e.g. various delay, cue, or reward durations) can be adjusted for one or all devices simultaneously. The new settings can be stored into each device’s onboard data storage to survive resets and power outages. Finally, each device is equipped with an interface accessible through a web browser which shows current parameter values and allows upload of a new program. After the initial upload, a cable connection is not necessary, and all protocol changes and code uploads are performed over the network. VLADISLAV devices also synchronise time, allowing sessions to be scheduled at a specific time of day. To our knowledge, VLADISLAV is the first operant HCM system with protocol templates, calendar time scheduling and full remote control and data acquisition. These features can be of particular interest in habitats with a high risk of contamination, such as germ-free or gnotobiotic facilities, to minimise the number of personnel entries.

Finally, the main design choice made to improve VLADISLAV’s robustness was implementing its entire functionality in an embedded microcontroller platform with no operating system or volatile memory. Therefore, unlike a Raspberry Pi or a PC, VLADISLAVs are resistant to corruption by power outages. Within one minute of restoring power, VLADISLAV reloads all saved protocol adjustments, becomes responsive to commands, and begins streaming data to as many redundant data storage devices as is required or reasonable. This level of robustness cannot be found in commercially available home cage operant systems and presents a key strength of the VLADISLAV architecture.

Further, more technical details may be found in Supplementary material 2 and the VLADISLAV repository (Virag, 2026). Additionally, VLADISLAV’s reduction of the necessary time investment is conservatively estimated in Supplementary material 3.

### What VLADISLAV Is, and What VLADISLAV Is Not

Today, new methods can be built upon a wealth of existing methods, platforms, and resources, a large majority of which didn’t exist only a few decades ago. Many of these are open source, available to researchers and companies alike (provided the terms of the projects’ licences are followed), reducing material costs, time, effort, and expertise required. The VLADISLAV system was developed by biomedical researchers supported by advice from electrical engineering (EE) experts. Furthermore, VLADISLAV is based on the MIROSLAV platform for locomotor activity monitoring, which was designed without any input from EE experts (Virag et al., 2025), much like many other devices developed by biomedical researchers (COST-TEATIME, n.d.; Kahnau et al., 2023). VLADISLAV was built in part to assess the feasibility of accomplishing the study aims with this approach and VLADISLAV, too, is now a resource available to everyone, including commercial entities. VLADISLAV’s successful implementation through transfer of open source platforms into the home cage shows that modern architectures should be readily available with negligible investments in research and development, and with few downsides compared to current, costly systems.

Importantly, VLADISLAV can be a usable alternative to existing systems with refinement, but has immediate value as an open source platform for reliable operant HCM. Its “copyleft” licencing allows replication, distribution, and all forms of modification by researchers and technicians comfortable with DIY electronics and commercial entities alike, while ensuring the platform and its subsequent developments remain open source.

Further development towards more user-friendly day-to-day use could be aimed at providing graphical interfaces to specify the initial network setup, protocol templates and parameters, and a web browser interface with dashboards for graphical real time monitoring (as seen in commercial Pallidus (Kravitz, 2022) and DVC (Tir et al., 2025a) systems). All of these potential benefits are also directly, inherently enabled by VLADISLAV’s networked architecture.

### MIROSLAV and VLADISLAV as a Manifesto for Modernising HCM System Architectures

The “black box” methodology of commercial scientific methods has long been criticised (Oellermann et al., 2022; Prager et al., 2019; Virag et al., 2021), but it may have even deeper ethical implications in HCM. No complex systems, including commercial ones, are free from unexpected issues, with practical advice and some descriptions of failures published by independent researchers and company affiliates (Grieco et al., 2021; Kahnau et al., 2022; Lipp et al., 2023; Mingrone et al., 2020; Tir et al., 2025b; Voikar & Gaburro, 2020). In case of severe failures where the experiment is in jeopardy, the responsibility lies with the researchers, but the outcome depends directly and solely on the company’s handling of the situation – including invested funds, work hours, and animal lives. Scientific data may be lost even if the company simply tends to the issue with a few days’ worth of delay due to the previously posited “hourglass” nature of animal experiments. Since the device is a black box, the researchers also cannot bring in external expertise to expediate the repair.

An earlier black box process, the design of a device, needs to be examined to answer questions of why the failure occurred and how likely is it to recur. The chance of failure, i.e. reliability, is primarily determined by the device’s underlying electronic architecture. If it cannot be examined, data on reliability and expected failure modes is indispensable, and commonly provided in other industries with devices in high-risk roles. While it is reasonable to expect researchers to test the device in their specific experimental setting upon purchasing it, stress testing and failure analyses require specialised electronic equipment and engineering expertise (Gandhi et al., 2019; Martin, 1999; Whitman, 2003). If any such analyses were conducted internally prior to marketing an HCM device, there is no requirement to disclose this information.

When designing or repairing a device, the company is under no obligation to provide detailed information on stress testing, discovered common failure modes, and estimated reliability regarding the system’s design, nor the cause, the solution, or verification process of a repair, if any. Importantly, there is no reason to doubt that companies conduct this work respecting the same ethical obligations which bind researchers. However, (mis)trust is a moot point – implementing and continuously reassessing precautions and safeties is prudent in any high-stakes situation where all risk is focused at a single point. Therefore, if a situation can occur where scientific data generation using public funds and animal lives fully depends on a single entity internally conducting a black box process, it is dangerous to eschew any scrutiny regardless of trust.

Following this principle, this study is, to our knowledge, the first direct attempt to open a discussion of reliability in HCM based on our work and available data. As a direct result of the lack of available information on reliability overall, and especially on HCM systems, the methodology of modelling reliability is coarse by necessity (limitations are further discussed in Supplementary material 2). Ultimately, even in the context of these limitations, the estimated difference of multiple orders of magnitude in reliability between PC-based and embedded architectures is extremely worrying. On the other hand, the question of feasibility and costs of such an architecture is answered by i) embedded networked architectures seeing widespread use in other industries for over a decade (Shehu Yalli et al., 2024), ii) VLADISLAV being implemented by researchers with no formal technical education, relying on the open source ecosystem, and iii) at a cost of about €1600 per 40 cage devices, including power supplies and Wi-Fi routers.

Purchasing support from companies guarantees expert system management while allowing researchers to invest their time and efforts elsewhere, which is a trade-off of immense importance. However, there is demonstrable room for progress in data loss prevention and reliability in state-of-the-art HCM. Keeping in mind the deep ethical implications, should we scrutinise our acceptance of obsolete architectures in commercial HCM more deeply and require more transparency? Can it be done without compromising the companies’ sustainability in a small market by examining open source business models?

With HCM rapidly evolving, tackling these questions now is crucial in ensuring safe and robust HCM methodology. In this debate, VLADISLAV is an example that we can do better – let’s discuss how.

## Conclusion

The worst possible outcome of data loss is the repetition of an experiment, wasting effort and, most importantly, animal lives. There is an ethical imperative to improve robustness in HCM, especially with very few commercial alternatives to HCM systems based on frail, inappropriate PC-based architectures which currently dominate the market. To better examine the problem, a networked embedded system was designed to conduct complex operant conditioning paradigms in the home cage – VLADISLAV. Through a pilot study, this work demonstrates the feasibility of utilising the open source ecosystem to build a reliable platform deployed across tens of cages simultaneously, with practical remote monitoring and batch control, at low cost and with little to no electronic engineering expertise. Indeed, such architectures have been widely used in other industries over the past decade to similarly remarkable effect. However, in commercial HCM, developments towards more reliable systems are scarce and closed processes dominate. Such a landscape has generally been accepted by the scientific community as necessary to keep the market of HCM systems sustainable.

In the context of this study’s findings, are more modern systems truly commercially infeasible? Are open processes truly at odds with commercial viability? If we had a choice, if they were open source, would we buy today’s systems at the same price if cheaper clones existed, just for expert support from the original developers? Realistically, is the current landscape of commercial HCM truly the only possible one? These questions are not rhetorical, and the answers are not implied – only by discussing these issues can we facilitate today’s use and tomorrow’s development of safe and reliable HCM methodology.

## Acknowledgements

The authors would like to thank laboratory technician Božica Hržan for supervising VLADISLAV devices and ensuring their continuous operation, as well as animal caretakers Miroslava Jerković, Tomislav Petanjek, and Anita Šantak for their patience while handling VLADISLAV-equipped cages.

The graphical abstract, Figure 1, the left panel of Figure 2, and Figure 3 were created in https://BioRender.com.

## Declarations

### Ethics Approval

All animal procedures were conducted in accordance with the Croatian Animal Protection Act (Class: 011-01/17-01/77, Reg no: 71-06-0/1-17-2), aligned with European Union legislation. All animal procedures were approved by the Croatian Ministry of Agriculture (EP 378/2022).

### Consent to Participate

Not applicable

### Consent for Publication

Not applicable

### Availability of Data and Materials

All data presented in the study and a full bill of materials to construct VLADISLAV can be found in the VLADISLAV repository (Virag, 2026) at the following URL: https://doi.org/10.5281/zenodo.18300841

Additional information can be found in Supplementary materials, as follows:

• **Supplementary material 1:**

Details on the Proof-of-Concept Study

• **Supplementary material 2:**

Hardware and Software Details, Network Security, Further Improvements

• **Supplementary material 3:**

Effects of VLADISLAV Architecture on Time Investment

## Code Availability

All VLADISLAV-related code, including Arduino/C++ embedded firmware, Python data acquisition scripts, Python data cleaning, preparation, and exploratory visualisation scripts, and R analysis scripts can be found in the VLADISLAV repository (Virag, 2026). URL: https://doi.org/10.5281/zenodo.18300841

## Authors’ Contributions

**DV**, **AMV**, and **JH** conceptualised the VLADISLAV platform. **DV** designed the VLADISLAV embedded circuit and infrastructure with critical input from **MC**. **DV** developed the VLADISLAV firmware with critical input from **JH** and **AMV**. **AMV** designed the VLADISLAV cage setup and optimised it for the experimental setting with critical input from **DV**. **AMV** produced all VLADISLAV devices with assistance from **DV**, **JH**, **ABP**, and **JOB**. **DV** performed data preparation and statistical analysis with critical input from **VT** and **JH**. **AMV** prepared Figures 1-3 with critical input from **DV**, **PK**, and **MŠP**. **DV** prepared Figures 4-7 with critical input from **JH**, **VT**, and **MŠP**. **PK** and **DV** empirically estimated time investment (methodology). In the Proof-of-Concept experiment, **DV**, **JH**, **ABP**, **AKn**, and **JOB** carried out the streptozotocin treatments. **DV** performed the behavioural testing. **DV** and **AMV** wrote the manuscript. **DV** created the graphical abstract. **AMV**, **JH**, **PK**, **ABP**, **AKn**, **AKr**, **LM**, **JOB**, **MC**, **VT**, and **MŠP** commented on the manuscript (including the graphical abstract and supplemental material) and provided critical feedback. **MŠP**, mentor of DV and the laboratory’s PI during this research, provided funding. Supervision was carried out by **JOB** and **MŠP**.

## Open Practices Statement

A version of this paper was submitted to bioRxiv. Experiments were not preregistered.

All code related to the manuscript, uploaded in the VLADISLAV repository (Virag, 2026), is licensed under GPLv3, a “copyleft” licence which ensures the code remains open-source, available to all for use, distribution, and modifications, provided that the derivative code is also licensed with GPLv3 or another, compatible “copyleft” license, ensuring the derivative code remains open-source as well.

---

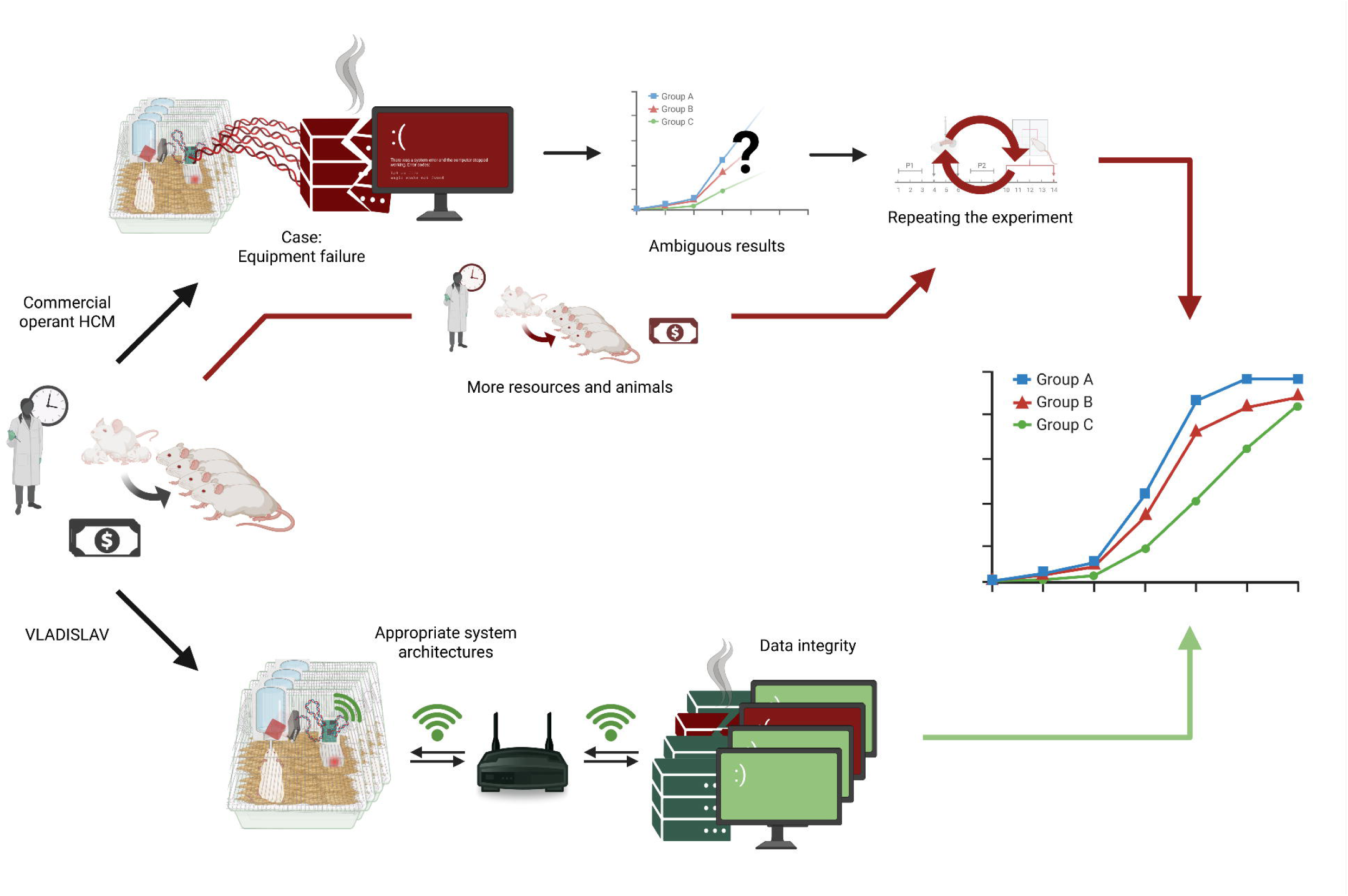

