## Supplementary material 1 for "Ethical Considerations of Mitigating Data Loss: VLADISLAV, a Manifesto for Reliable Home Cage Systems"

*Supplementary material 1* **Details on the Proof-of-Concept Study**

***Why Home Cage, and Why Not Existing Systems?***

While designing the study, which is part of a larger project, several issues needed to be considered to implement an operant test.

**Training duration.** Firstly, conventional, out-of-cage paradigms require habituation and training periods of around one month, which cannot offer insight into early treatment effects. Furthermore, at least 1 hour/day/animal of required training makes the test prohibitively impractical and/or expensive, especially in 30-40-animal studies common in a 2x2 factorial experimental design. Parallelised testing using multiple operant chambers would expend available space before reaching time efficiency. **Solution:** A home cage system.

**Reward choice.** Secondly, both conventional and home cage operant systems are usually built for dispensing sweet, sugary pellets or water solutions as rewards. Given the effects of streptozotocin in insulin-related metabolic pathways (Knezovic et al., 2017, 2023) and the beneficial effects of oral galactose on the model’s pathology (Knezovic et al., 2018), modifying the diet could introduce a potential confounder. Optimally, drinking water would be used with *ad libitum* access outside of testing sessions, or, at worst, with mild water deprivation (norecopa.no, 2023). These requirements narrow the choice to setups with gated access to a bottle, excluding pellet dispensers and syringe pump liquid dispensers. **Solution:** A system with gated access to a standard bottle with tap water.

***Observations on the Necessity of Water Deprivation***

Two animals reused from other experiments were exposed to the device to ensure errorless device operation for the full experiment according to the Reuse principle of the 3Rs. Under continuous remote video supervision, we observed both animals learning to operate the VLADISLAV apparatus within hours, without water deprivation beforehand (data not shown). Though these findings aren’t rigorously evaluated, when put in the context of some alternate protocols reported in the literature (Augier et al., 2017; Reinagel, 2018), it is likely that the necessity of water deprivation for operant conditioning should be reconsidered, especially in the home cage context, however, a systematic evaluation is required.

***Single Housing in Home Cage Operant Conditioning***

Since rats are social animals, single housing in social isolation has a profound effect (Begni et al., 2020) and is an important limitation of this study, deemed necessary due to two reasons. Firstly, similar to some other operant devices, VLADISLAV cannot monitor which animal solves a task, and cannot ensure that the same animal will consume the reward in its current setup. Secondly, single housing was required for precise individual dosing of treatment via drinking water as part of the full study from which the presented data is subset. As social housing is an important component of laboratory rodents’ welfare, alternative approaches are intensely considered for future studies. These issues are commonly associated with home cage operant conditioning (HCOC) devices, such as VLADISLAV and FED3 (Matikainen-Ankney et al., 2021), but arise due to a separate issue of access control. Home cage access control systems are a separate challenge in electrical engineering, similar to HCOC itself.

Few systems exist which integrate HCOC with access control, with the TSE IntelliCage likely being the most well-known. This specific system’s architecture is PC-dependent, with the architecture’s reliability and ease of use compared in this work to VLADISLAV’s contemporary networked architecture, seen commonly in other fields for more than a decade.

A possible way to approach both issues is to connect a “living” cage, which would house multiple animals, with a testing/treatment chamber using Radio Frequency Identification (RFID) gating. Such gating systems have been developed and published by Winter & Schaefers (Winter & Schaefers, 2011) and Mei et al. (Mei et al., 2020). An integrated system would detect implanted RFID tags and only let one animal through to the chamber. The chamber would contain a VLADISLAV gating access to each animal’s bottle, and, if the trial is solved correctly, only access to the appropriate bottle would be allowed based on the animal’s individual tag ID retrieved from the gate. VLADISLAV’s open architecture and built-in precise time synchronisation will facilitate its integration with other systems. Precise timekeeping is important when integrating data from multiple systems in general, which will be discussed in detail in a book on home cage monitoring systems, which will be published in 2026 by Springer Nature, edited by Stefano Gaburro and Silvia Mandillo.

***Blinding***

This work utilises a subset of experimental data to demonstrate VLADISLAV’s feasibility. Two additional treatment groups were present in the full experiment (manuscript in preparation). Due to the precise method the second treatment’s daily application, while blinding was possible, it was determined during the study design phase not to be feasible given the available resources and personnel. In the context of this limitation, the findings should serve to navigate further exploration of the phenomena observed, including replication in a blinded design.
