## Supplementary material 2 for "Ethical Considerations of Mitigating Data Loss: VLADISLAV, a Manifesto for Reliable Home Cage Systems"

Supplementary material 2
VLADISLAV: Hardware and Software Details, Network Security, Further Improvement

### Build Details

From the hardware side, VLADISLAV is built to be a simple, flexible device with three main electronic parts: i) the microcontroller board, with an RGB LED, used as a neutral cue, ii) a servo, gating access to water, and iii) a beam-break sensor to detect nosepokes. Mechanical components (gate, nosepoke opening, bottle holder...) can be customised depending on the experimental requirements, cage type, and accessible tooling and materials. An electronic circuit connects the components, with a schematic and a full bill of materials shared in the VLADISLAV repository (Virag, 2026). The total cost of 40 devices is around €1600. To accommodate a wide variety of rodent enclosures, the number of components is kept at a minimum. The mechanical parts can be constructed using off-the-shelf parts available in most hardware or hobby supply stores (nuts and bolts, aluminium profiles, plexiglass or polystyrene panels) and processed using a saw and a drill, or 3D-printed. Though 3D printing is a common option, especially with mice, we opted for a combined plexiglass-aluminium build with all parts internal to the cage fashioned out of metal for endurance. It is possible that the increased exploration up to the point of destruction of plastic material is exacerbated by the lack of cage enrichment.

### Prepare-a-VLAD

The notebook has three main, sequential objectives:

1. Loading data from textual logs into a *Pandas* data frame with the appropriate animal and group labels,
2. Supplementing the data frame with experimental design parameters, and filtering based on events logged in experimental notes,
3. Exploring the data using heatmaps and performing additional filtering.

The final step involves raw data examination on a per-subject basis primarily to discover unexpected device operation and failures and filter the data accordingly, but to also explore novel, unexpected behavioural patterns as the first step in exploratory data analysis.

### Internet Access And Network Hardening

To access devices via Internet, a Virtual Private Network (VPN) can be used. To harden the network, all other Internet access can be prohibited, except VPN traffic. However, networking does not necessarily imply Internet access, and setting up an isolated Local Area Network (LAN) without Internet access is a good way to decrease the attack surface. To prevent unauthorised access, newer Wi-Fi security protocols can be used (e.g. WPA3), or the devices could be switched to a wired Ethernet connection completely, enabled by add-on peripheral boards. Within the network, the Message Queuing Telemetry Transport (MQTT) protocol supports TLS encryption to prevent man-in-the-middle attacks. The specific measures will depend on the facility and specific circumstances, but a network of VLADISLAVs, or any other HCM devices, need not be inherently unsafe.

However, the architecture can still be made more robust and determining the cost-benefit relationship of implementing further measures would be of practical use. Importantly, some institutions may object to networking. It should be noted that state-of-the-art security features can be implemented with ESP32-based devices, and the network used for redundant data acquisition and storage can be completely isolated from both the institution’s network, and the Internet, by simply not connecting the router or any of the recording devices to any other networks. Furthermore, instead of Wi-Fi, other types of networks can also be used for this purpose, such as LoRa, ZigBee, Bluetooth.

### The Curious Case of Calculating Reliability

The calculated difference is based on reliability curves of PCs and mean time to failure (MTTF) of routers. MTTF is an imperfect measure, since, unlike the reliability curve, it does not describe the distribution – individual devices in various circumstances can fail very quickly or greatly exceed the stated MTTF. The distribution of device failure usually follows a “bathtub” curve (Ohring, 1995). Furthermore, it is likely that a PC dedicated to running HCM software would experience less failures and have a longer mean time to failure (MTTF), but an estimated 200-fold difference is worrying even when permitting a large margin for error. Lastly, VLADISLAV’s mode of failure does not allow for retrograde data loss, which is possible with a PC-based system.

### The Router Is Still a Single Point of Failure, Isn’t It?

In this work, the desktop PC’s reliability is criticised and positioning it as a single point of failure (a component critical for entire system operation) is advised against. In the VLADISLAV setup, the single point of failure still exists, but it has been shifted to the Wi-Fi router. The router is an embedded system made for prolonged operation with minimal downtime, and thus more robust than a desktop PC, as demonstrated in the study, but additional measures are possible. The ESP32-S2 microcontroller used in the study contains 4 megabytes of flash memory, used to store protocol templates and parameters. This memory can be exposed as a storage device on USB connection to a computer, and could be utilised as a circular buffer for storing most recent messages. The messages could be retrieved via USB in case of prolonged network outage. If a larger buffer would be required, the FED3 (Matikainen-Ankney et al., 2021) approach could be employed by extending VLADISLAV with memory card storage.

### Scaling Non-Networked Architectures

With 40 simultaneous devices, the discussed concerns regarding desktop PCs would be exacerbated by needing to connect many devices to one PC, whereas using many PCs to acquire relatively simple data is not cost- nor energy-effective, and would possibly produce too much heat and noise. The FED3 (Matikainen-Ankney et al., 2021) is economic in terms of cost and space but would require physically manipulating every device to retrieve data or load a new protocol via its memory card. Furthermore, it is built to dispense sucrose pellets, which cannot be used in our disease model due to its metabolic component.
