## Supplementary material 3 for "Ethical Considerations of Mitigating Data Loss: VLADISLAV, a Manifesto for Reliable Home Cage Systems"

*Supplementary material 3* **Effects of VLADISLAV Architecture on Time Investment**

The architectural differences described in the main text have a direct impact on the amount of time which needs to be spent on the two devices throughout an experiment. Specifically, due to VLADISLAV’s design, data retrieval does not exist as a task requiring human intervention. To reduce bias, though the estimates are empirical, all task durations (listed in column “Task duration” of Table S1) were estimated to be the same for both devices with the exception of data retrieval which does not exist as a task for VLADISLAV. The duration of other tasks was estimated regardless to be able to put time savings into context of a total time investment estimate.

Furthermore, since data retrieval is automatic and continuous for VLADISLAV, the time investment needed to be assessed in reasonably similar circumstances for the PC-controlled device – therefore, it was decided to fix the frequency of data retrieval for the PC-based device to “daily”. Thus, to more accurately assess time investment, the confounding difference in data loss risk is minimised between the devices and analysed separately, as described in the main text.

The time investment was assessed two hypothetical workers (researchers, technicians, caretakers, or in other roles) with different roles and responsibilities in the experiment, but directly maintaining the device’s operation day-to-day. In a “high workload” scenario, the worker enters the habitat every day by default due to other duties (to perform a mandatory daily check of the animals), and uses the occasion to perform a visual inspection of the device and retrieve data, if applicable. In a “low workload” scenario, the worker does not need to perform daily checks (the task is delegated to someone else), so they enter the habitat only once per week, and more frequently if required for the maintenance of the device – entrance is assumed to be required daily for the commercial device to perform data retrieval, and weekly for VLADISLAV to perform an overall device inspection.

Despite the coarseness and limited practical value of the estimate, it is the first such comparison in the literature to the authors’ knowledge, intended to give a sense of scale and encourage future, more rigorous studies directly comparing multiple systems.

| **Table S1**  *Empirical Estimates of Home Cage Operant Device-Related Tasks in a Typical Experiment* | | | |
| --- | --- | --- | --- |
| **Task** | **Task frequency over the course of an experiment** | | **Task duration** |
|  | High workload scenario | Low workload scenario |  |
| Design protocols | One time | One time | 45 minutes |
| Load a protocol | Once every 2 weeks | Once every 2 weeks | 5 minutes |
| Habitat entry | Every day | **VLADISLAV:** Once every week  **Commercial device:** Every day | 15 minutes |
| Retrieve data | **VLADISLAV:** N/A **Commercial device:** Every day | **VLADISLAV:** N/A **Commercial device:** Every day | 5 minutes |
| Data inspection | Every day | Every day | 10 minutes |
| Visual inspection | Every day | Once every week | 10 minutes |
| System cleaning and restart | Once every week | Once every week | 60 minutes |
| *Note.* Estimates given apply to both VLADISLAV and a typical PC-controlled commercial device, unless otherwise specified. The “Retrieve data” task does not exist in the VLADISLAV workflow by design, as described in the text. | | | |

Figure S1 shows estimated cumulative time spent on maintenance tasks for high and low time investment scenarios for both devices (as described in Materials and Methods) in the panel on the left. The panel on the right shows the difference, corresponding to time saved with VLADISLAV depending on experiment duration.

**Figure S1**

*Estimated Cumulative Time Spent on Device Maintenance*

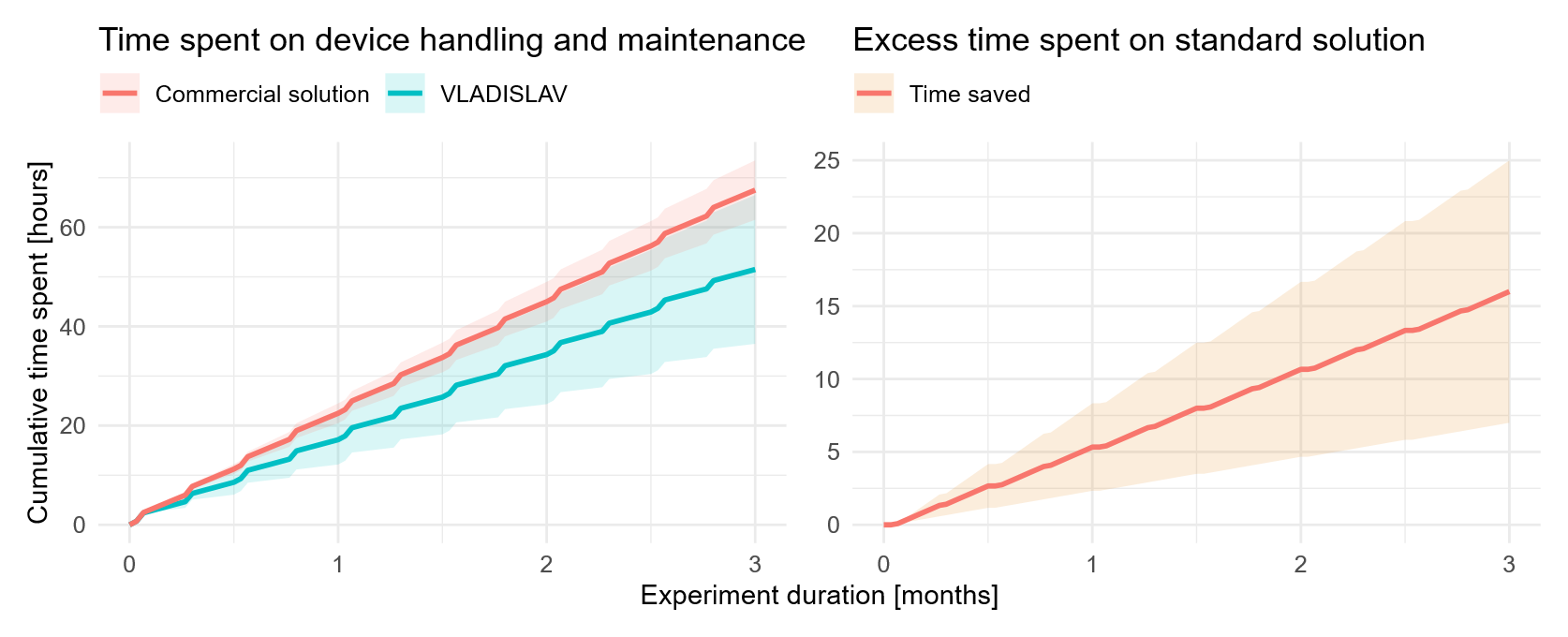

*Note.* Per-device estimates are shown on the left. The lower end of each ribbon corresponds to estimates for the low time investment scenario, the upper end to estimates for the high time investment scenario, and the line shows the calculated mean of the two scenarios. The plot on the right shows the difference between the two devices, corresponding to cumulative time saved with VLADISLAV over the duration of an experiment.
